# Autophosphorylation of the Tousled-like kinases TLK1 and TLK2 regulates recruitment to damaged chromatin via PCNA interaction

**DOI:** 10.1101/2024.04.22.590659

**Authors:** Kirk L. West, Natasha Kreiling, Kevin D. Raney, Gargi Ghosal, Justin W Leung

## Abstract

Tousled-like kinases 1 and 2 (TLK1 and 2) are cell cycle-regulated serine/threonine kinases that are involved in multiple biological processes. Mutation of TLK1 and 2 confer neurodegenerative diseases. Recent studies demonstrate that TLK1 and 2 are involved in DNA repair. However, there is no direct evidence that TLK1 and 2 function at DNA damage sites. Here, we show that both TLK1 and TLK2 are hyper-autophosphorylated at their N-termini, at least in part, mediated by their homo-or hetero-dimerization. We found that TLK1 and 2 hyper-autophosphorylation suppresses their recruitment to damaged chromatin. Furthermore, both TLK1 and 2 associate with PCNA specifically through their evolutionarily conserved non-canonical PCNA-interacting protein (PIP) box at the N-terminus, and mutation of the PIP-box abolishes their recruitment to DNA damage sites. Mechanistically, the TLK1 and 2 hyper-autophosphorylation masks the PIP-box and negatively regulates their recruitment to the DNA damage site. Overall, our study dissects the detailed genetic regulation of TLK1 and 2 at damaged chromatin, which provides important insights into their emerging roles in DNA repair.

## Introduction

Serine/threonine kinases, Tousled-like kinases 1 and 2 (TLK1 and 2), are implicated in DNA repair, immune response, chromatin structure, transcription, DNA replication, checkpoint control, and cell morphology (1–19). Mutations of TLK1 and 2 are associated with neurodevelopmental disorder (9, 20–24).

A growing body of evidence shows that TLK1 and 2 are key DNA repair proteins. The activity of TLK is cell cycle-regulated peaking during S-phase progression (12, 25) and transiently inhibited after DNA double-strand breaks (DSBs) (26). The DNA damage-induced inhibition is regulated by ATM and Chk1 (26, 27).

Although many TLK substrates have been reported, the H3/H4 histone chaperones, anti-silencing function-1 (ASF1A and ASF1B) (8, 28, 29), involved in chromatin assembly and DNA repair via CAF1 and HIRA, are the most well characterized (30–35). Besides ASF1A and ASF1B, several DNA repair proteins, including NEK1, RAD9, and RAD54 (16, 36–39), were identified as TLK substrates indicating an important role of TLKs in maintaining genome stability (3, 39, 40). Both TLK1 and TLK2 are enriched at nascent DNA at replication fork (41). However, with their previously reported functions in stabilizing replication fork and involvement in homologous recombination (1, 39), there is no direct evidence that TLKs are localized at the sites of DNA damage.

Here, we investigate the molecular mechanism by which TLK1 and 2 are regulated at damaged chromatin. Our data suggest that TLK1 and 2 are recruited to laser-induced microirradiation and their recruitment is inhibited by their kinase activity. Using proteomic analysis, systematic mutagenesis, and biochemical assays, we have identified autophosphorylation sites at the N-terminus of TLK1 and 2. Importantly, auto-hyperphosphorylation of TLK1 and 2 inhibits their recruitment to damaged chromatin. Mechanistically, recruitment of TLK 1 and 2 is mediated by PCNA through their putative PCNA interacting protein box (PIP-box) that is masked by the surrounding phosphorylated residues. Our study provides novel molecular insights into TLK activity that regulates their localization to damaged chromatin.

## Materials and methods

### Cell culture

HEK293T and U2OS cell lines were obtained from American Type Culture Collection. U2OS H2AX knockout (KO), RNF168 KO, 53BP1 KO, and LC8 KO cells were generated using CRISPR/Cas9 technology as previously described (42). All cells were cultured in Dulbecco’s modified Eagle’s medium (DMEM) with 10% fetal bovine serum supplemented with 100 U/mL penicillin and 100 µg/mL streptomycin at 37°C with 5% CO_2_. Polyethylenimine (PEI) (Polysciences, 24765-1) was used for transfection. PEI and plasmids were mixed in a 3 µl: 1 µg ratio in OPTI-MEM, incubated for 10 minutes and added to the medium for 6 h.

### Antibodies

Primary antibodies Flag M2 (Sigma, F1804), TLK1 (Cell Signaling, 4125s), TLK2 (Bethyl, A301-257A), Myc (Santa Cruz, sc-40), GFP (Invitrogen, A11122), β-Tubulin (Abcam, ab6046), Histone H4 (Cell Signaling, 2935S), LC8 (Abcam, ab51603), Thiophosphate ester (Abcam, ab92570), Phosphoserine (Millipore, AB1603), Phosphothreonine (New England Biolabs, 9386S), PCNA (Abcam, ab92552) and secondary antibodies Goat Anti-Mouse IgG-HRP (Jackson ImmunoResearch,111-035-144) and Goat Anti-Rabbit IgG-HRP (Jackson ImmunoResearch, 115-035-166) secondary antibodies were used for western blotting analysis.

### Plasmids

Human TLK1 pDONR221 was purchased from the Harvard PlasmID Database (Plasmid #HsCD00042921). Human TLK2 pDONR223 was a gift from William Hahn & David Root (Addgene: Plasmid #23629). Human LC8 expression vectors and its mutants were described previously (42). pDONR vectors were subcloned into indicated gateway-compatible destination vectors using Gateway LR Clonase II (Invitrogen, 11791020). TLK1 and TLK2 point mutants were generated by site-directed mutagenesis reactions using the NEB Q5 site-directed mutagenesis kit (NEB, E0554S). Primers are listed in Supplementary Table 1.

### Tandem affinity purification

Tandem affinity purification (TAP) was carried out as described previously (43). Briefly, S-protein-2xFLAG-Streptavidin binding peptide (SFB)-tagged TLK1, SFB-TLK1 1-240, SFB-TLK1 D607A, SFB-TLK1 133-208, SFB-TLK1 133-208 Y149A F150A, or SFB-TLK2 were transfected into HEK293T cells. Cells were harvested 24 hours after transfection using NETN buffer (150 mM NaCl, 0.5 mM EDTA, 20 mM Tris-HCl pH 8.0, 0.5% NP-40) supplemented with 2 µg/mL aprotinin (Thermo, AAJ60237MB) and 5 µg/mL pepstatin A (Thermo, PI78432) at 4°C for 20 minutes. Lysates were centrifuged at 9,000 x g, 4°C for 20 minutes to yield supernatant as soluble fraction. The pellet was washed 1x with PBS and centrifuged at 9,000 x g, 4°C for 5 minutes and lysed with chromatin extraction buffer (NETN buffer, 10 mM MgCl_2_, and Turbonuclease, Accelagen, N0103M) at 4°C for 1 hour followed by centrifugation at 9,000 x g, 4°C for 20 minutes to obtain chromatin fraction. Both soluble and chromatin fractions were incubated with streptavidin sepharose (200 μl) (GE Healthcare, GE17-5113-01) at 4°C for 1 hour followed by washing with NETN buffer three times. The protein complexes were eluted with 2 mg/mL biotin at 4°C for 1 hour. The eluents were then incubated with S-protein agarose (EMD Millipore, 69704-3) overnight at 4°C, washed three times with NETN buffer, and eluted in 1x Laemmli buffer.

### Mass spectrometric analysis of protein-protein interactions

Protein complexes obtained from TAP were processed by in-gel trypsin digestion and analyzed by tandem mass spectrometry (44). Briefly, protein samples were electrophoresed on a 4-20% tris-glycine gel for 5 minutes at 300V. The gel was stained with GelCode Safe (Thermo Scientific, 24594) according to the manufacturer’s instructions. The excised gel band was destained three times with 50 mM ammonium bicarbonate (Sigma-Aldrich, A6141) and 50% methanol (Fisher, A456) for 1 hour each. Each sample was dehydrated by acetonitrile (Fisher, A955). Gel bands were reduced with 10 mM Tris(2-carboxyethyl)phosphine) (TCEP) (Pierce, 20490) for 30 minutes at 37°C. Free thiol groups were derivatized by 50 mM Iodoacetamide (Sigma-Aldrich, A3221) for 1 hour at room temperature. Gel bands were washed by alternating 100 mM ammonium bicarbonate and acetonitrile three times, followed by dehydration with acetonitrile. Samples were digested by sequencing grade trypsin (Promega, V5111) in 50 mM ammonium bicarbonate at 37°C overnight. The samples were then acidified by 0.5% formic acid (Fisher, A117) to a final concentration of 0.1% and were desalted by Water’s C18 SepPak (Waters, WAT023590), dried by SpeedVac and resuspended in 0.1% formic acid. The samples were then injected into an in-line, reverse phase, 150 x 0.075 mm column packed with XSelect CSH C18 2.5µM resin (Waters, 186006103) using an UltiMate 3000 RSLCnano. Peptides were eluted into an Orbitrap Fusion Tribrid mass spectrometer following a 100-min gradient of 97:3 Buffer A (0.1% formic acid and 0.5% acetonitrile) to buffer B (0.1% formic acid and 99.9% acetonitrile) to 67:33 A:B. Ionization of eluted peptides was performed by electrospray ionization at 2.2 kV. Precursor ions were collected in the Orbitrap at 240,000 resolution for a MS^1^ range of 375 – 1,500 m/z. Data-dependent acquisition was used to select the top 10 precursor ions for fragmentation by higher-energy collisional dissociation with normalized collision energy between 28.0 and 31.0. Fragment ions between 200 – 1,400 m/z were measured at MS^2^ in the ion trap.

Proteins were identified using Mascot (Matrix Sciences) to search the *Homo sapiens* UniProtKB database. Search conditions were set to a parent ion tolerance of 3 ppm and a fragment ion tolerance of 0.5 Da. Carbamidomethylation of cysteine was selected for fixed modifications. Oxidation of methionine, acetylation of peptide N-termini, and phosphorylation of serine, threonine, and tyrosine were selected as variable modifications. Scaffold (Proteome Software) was used to validate protein and peptide identifications. Proteins and peptides were identified if they exhibited less than 1.0% false discovery by Scaffold’s local false discovery algorithm.

### TMT11plex labeling and phosphopeptide analysis

Peptide labeling and phosphopeptide enrichment were carried out as previously described (45). Briefly, HEK293T cells were transfected with either SFB-TLK1, SFB-TLK2, siRNA targeting TLK1, TLK2, or TLK1 and TLK2. Cells were harvested twenty-four hours after transfection, four replicates of each transfected or control non-transfected cells were lysed in 2% SDS and 100 mM Tris-HCl, pH 7.6 supplemented with fresh protease (Pierce, A32963) and phosphatase inhibitors (Pierce, A32957). Protein concentrations were measured by BCA protein assay (Pierce, 23225). 300 µg of protein from each sample was reduced by TCEP, alkylated by iodoacetamide, and purified via chloroform/methanol extraction. Samples were then trypsinized with a protein: trypsin ratio of 50:1 overnight at 37°C followed by quenching with 0.5% formic acid to a final concentration of 0.1%. Samples were salted by Waters C18 SepPaks and eluted peptides were dried using a SpeedVac. 120µg of peptides were labeled using a TMT10plex isobaric label reagent set with the addition of the TMT11-131C label (Thermo, A34808). Labeled peptides were dried by speed-vac and resuspended in phosphopeptides binding/wash buffer from the High-Select TiO_2_ phosphopeptide enrichment kit (Pierce, A32993). Samples were fractionated using a 100 x 1.0-mm Acquity BEH C18 column (Waters) by an Ulitmate 3000 UHPLC system (Thermo) over a 40 minute gradient following ratio of 99:1 to 60:40 basic buffer A:B (buffer B: 10 mM ammonium hydroxide, 99.9% acetonitrile, pH 10) into 36 fractions. Multinotch MS^3^ was performed as previously described(46). Data analysis was performed using Maxquant (Max Planck Institute)(45, 47). The *Homo sapien* UniprotKB database was used for peptide searching, intensities were used to determine significantly enriched phosphopeptides by the ProteoViz R scripts with the R package limma (48, 49). Heatmaps were created in GraphPad Prism 10 (GraphPad).

### Streptavidin pulldown

HEK293T cells were transfected with SFB-expression vectors as indicated and harvested by addition of trypsin, washed with PBS, and then lysed by addition of NETN buffer with Turbonuclease for 1 hour at 4°C. Lysates were centrifuged at 14,000 x g, 4°C for 20 minutes. Supernatants were incubated with streptavidin sepharose beads for 1 hour at 4°C with rotation and washed three times with NETN buffer. Protein complexes were eluted 1x Laemmli buffer for Western blotting analysis.

### Chromatin fractionation

HEK293T cells were transfected with the SFB-expressing vectors. Cells were harvested NETN buffer supplemented with aprotinin and pepstatin A. Whole cell extracts were obtained from the direct lysate. The remaining lysate was centrifuged at 14,000xg, 4°C for 20 minutes. The supernatants were collected for the soluble fraction and the insoluble chromatin pellet was washed twice with PBS. The pellets were extracted using 0.2N hydrochloric acid for 15 minutes as the chromatin fraction and neutralized by equal volume 1M Tris pH 8.0.

### Western blotting

Samples were resolved by SDS-PAGE, transferred to PVDF membranes, blocked with 5% non-fat milk, and incubated with the indicated primary antibodies overnight at 4°C. The blots were washed for three times with 1X Tris-Buffered Saline with 0.1% Tween 20 (TBST). The membranes were incubated at room temperature for 1 hour with HRP-conjugated secondary antibodies. The blots were then washed three times with TBST and were developed using Clarity ECL (BioRad, Cat #1705061). Images were captured using a BioRad ChemiDoc MP.

### Laser-induced micro-irradiation

Cells were seeded on 35 mm glass bottom dishes (Cellvis, D35-20-1.5-N) and transfected with GFP-tagged proteins as indicated. Laser-induced microirradiation was performed using a Nikon Ti2 inverted fluorescent microscope and C2+ confocal system with an Okolab live cell imaging chamber attached to a gas-mixer maintained at a temperature of 37°C and 5% CO_2_. The 405 nm laser of the C2+ confocal system was used to irradiate each cell at the indicated region. Images were captured in 30-second intervals after damage and the fluorescent intensity of GFP-tagged protein recruitment to the damaged region was quantified. The intensity of the recruitment was normalized by subtracting the background intensity of an undamaged area within the same cell at the same time point.

### Kinase assays

Kinase assays were performed as previously described (50). Briefly, SFB-tagged proteins were expressed in HEK293T cells for 48 hours. Cells were lysed in NETN buffer with protease inhibitors, 10 mM MgCl_2_, and Turbonuclease for 1 hour at 4°C. Samples were centrifuged at 14,000xg, 4°C for 20 minutes. Supernatants were incubated with S-protein beads for 1 hour at 4°C. Beads were washed for three times with NETN, twice with kinase wash buffer (40 mM HEPES pH 7.5, 250 mM NaCl), and once with kinase reaction buffer (30 mM HEPES pH 7.5, 50 mM potassium acetate, 5 mM MgCl_2_). Equal volumes of SFB-tagged proteins in kinase reaction buffer and 500 µM ATP-γ-S (Abcam, ab138911) were mixed and incubated at 30°C for 30 minutes. ATP-γ-S was alkylated by p-Nitrobenzyl mesylate (PNBM, Abcam, ab138910) to a final concentration of 2.5 mM and incubated at room temperature for 1 hour. Alkylation was terminated using 2x Laemmli buffer. Phosphorylation was detected after Western blotting by an anti-thiophosphate ester antibody.

### Lambda protein phosphatase assay

HEK293T cells were transfected with SFB-expressing vectors. 24 hours post-transfection, cells were harvested in NETN buffer with protease inhibitors, 10 mM MgCl_2_, and Turbonuclease for 1 hour at 4°C. SFB-tagged proteins were purified using S-protein beads and washed for three times with NETN buffer. SFB-tagged proteins were aliquoted into a fresh 1.5 mL tube containing 1x NEBuffer for Protein MetalloPhosphatases (PMP) supplemented with 1 mM MnCl_2_, and 400 units of Lambda protein phosphatase (λpp) (New England Biolabs, P0753S). The reactions were incubated at 30°C for 30 minutes and terminated by an equal volume of 2x Laemmli buffer.

### Statistical analysis

Laser-induced micro-irradiation experiments represent the mean ± SEM of N ≥ 10 cells in each condition unless otherwise specified. Unpaired, two-tailed Student’s *t-*tests and one-way ANOVA followed by Dunnett’s multiple comparison test were performed in GraphPad Prism 10. Statistical significance corresponds to P < 0.05.

## Results

### TLK1 and TLK2 form homo- and heterodimers

Recent studies revealed that LC8 (also known as DYNLL1) is a DNA repair protein that functions through its interactions with 53BP1, a key regulator of DSB pathway choice (42, 51–53). Previously, we demonstrated the binding of LC8 to numerous DNA repair proteins, including CHD4, ZMYM2, and ZMYM3. Consistent with previous reports (54), we observed TLK1 and TLK2 were pulled down as LC8 interactors in our LC8 proteomic analysis (7, 42). Subsequent tandem affinity purification-coupled mass spectrometry of TLK1 and TLK2 revealed that TLK1 and TLK2 are each other’s primary interactors followed by LC8 (**Supplementary Figure 1A**).

In order to map the interactions between TLK1/2 and LC8, we systematically mutated the known domains of TLK1 and 2 (**Figure 1A-B**) and performed pulldown assays (**Figure 1C-E**). Consistent with previous work, we found that TLK1 and TLK2, both form homodimers and heterodimers through their coiled-coil 1 domain (**Figure 1C-E**) (54, 55). We identified the LC8 anchoring motifs on TLK1 (SFKI**<u>IQT</u>**D) and TLK2 (QHEQ**<u>TQS</u>**D) (**Supplementary Figure 1B-H**) as direct interaction sites for the LC8 canonical binding groove. However, LC8 was not involved in TLK1 and TLK2 dimerization (**Figure 1C-E**) despite the well-characterized function of LC8 in promoting the dimerization of its interacting partners, including other kinases such as NEK9 (42, 52, 56).

**Figure 1.**
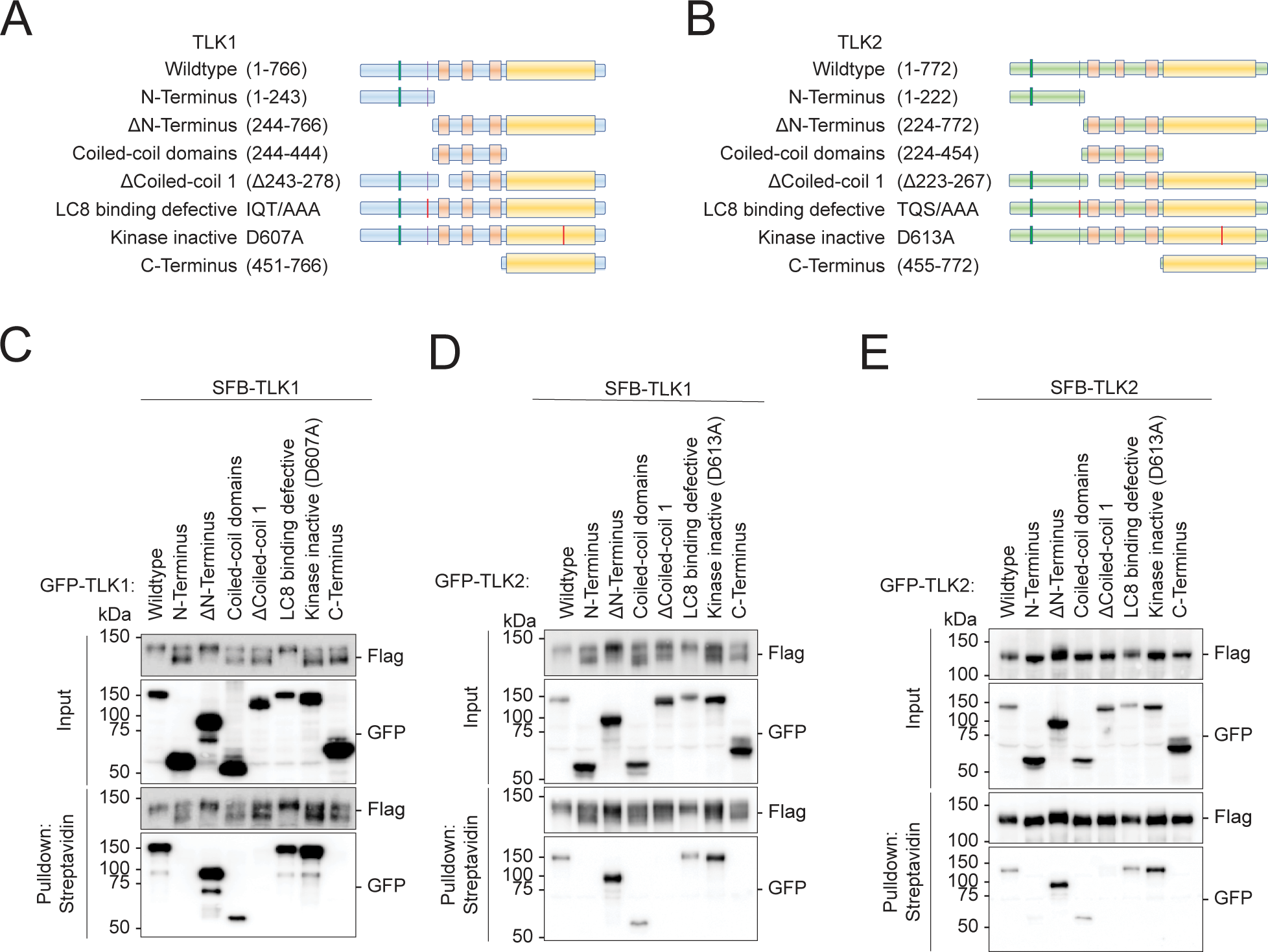
Tousled-like kinases form dimers and drive autophosphorylation. **(A-B)** Schematic illustrations of TLK1 and TLK2 constructs for dimerization domain mapping. **(C-E)** TLK dimerization requires an intact coiled-coil domain. HEK293T cells were co-transfected with **(C)** SFB-TLK1 wildtype and the indicated GFP-TLK1 expression vectors, **(D)** SFB-TLK1 wildtype and GFP-TLK2 plasmids, **(E)** SFB-TLK2 and GFP-TLK2 plasmids, and pulldown experiment followed by Western blotting analysis using FLAG and GFP antibodies.

### Dimerization of TLK1 and TLK2 drives autophosphorylation of the N-terminus

In the pulldown assay, we observed a pronounced differential migration rate for TLK1, and a modest difference for TLK2, when co-expressed with different mutants (**Figure 1C-E**). More specifically, SFB-TLK1 migrated slower when co-expressed with wildtype, ΔN-terminus, and LC8-binding defective mutants. Co-expression with the non-interacting (N-terminus, coiled-coil domain, Δcoiled-coil1, and C-terminus) or the catalytically inactive D607A mutants revealed two prominent forms: a faster and slower migrating TLK1 (**Figure 1C and D**). Interestingly, co-expression of GFP-TLK1 and 2 wildtype, ΔN-terminus and LC8-binding defective mutants shifted the SFB-TLK1 D607A to the slower migrating form (**Supplementary Figure 2A-B**), while we only observed the faster migrating bottom band for SFB-TLK1 D607A upon co-expression with non-interacting and catalytic inactive mutants (**Supplementary Figure 2A-B**). Consistently, a TLK2 catalytically inactive mutant did not show a drastic shift in size when co-expressed with wildtype or a number of different mutants (**Supplementary Figure 2C**). The mobility shift was dependent on the presence of the active form of TLK1. We only observed a faster migrating band when we co-expressed SFB-TLK1 D607A and GFP-TLK1 D607A (**Figure 2A**). Titrating the wildtype plasmid with GFP-TLK1 D607 mutant construct, we observed a mobility shift to a slower migrating band (**Figure 2A**). The mobility shift correlated with the presence of TLK1 activity, indicating that the upper band of TLK1 is the phosphorylated form.

**Figure 2.**
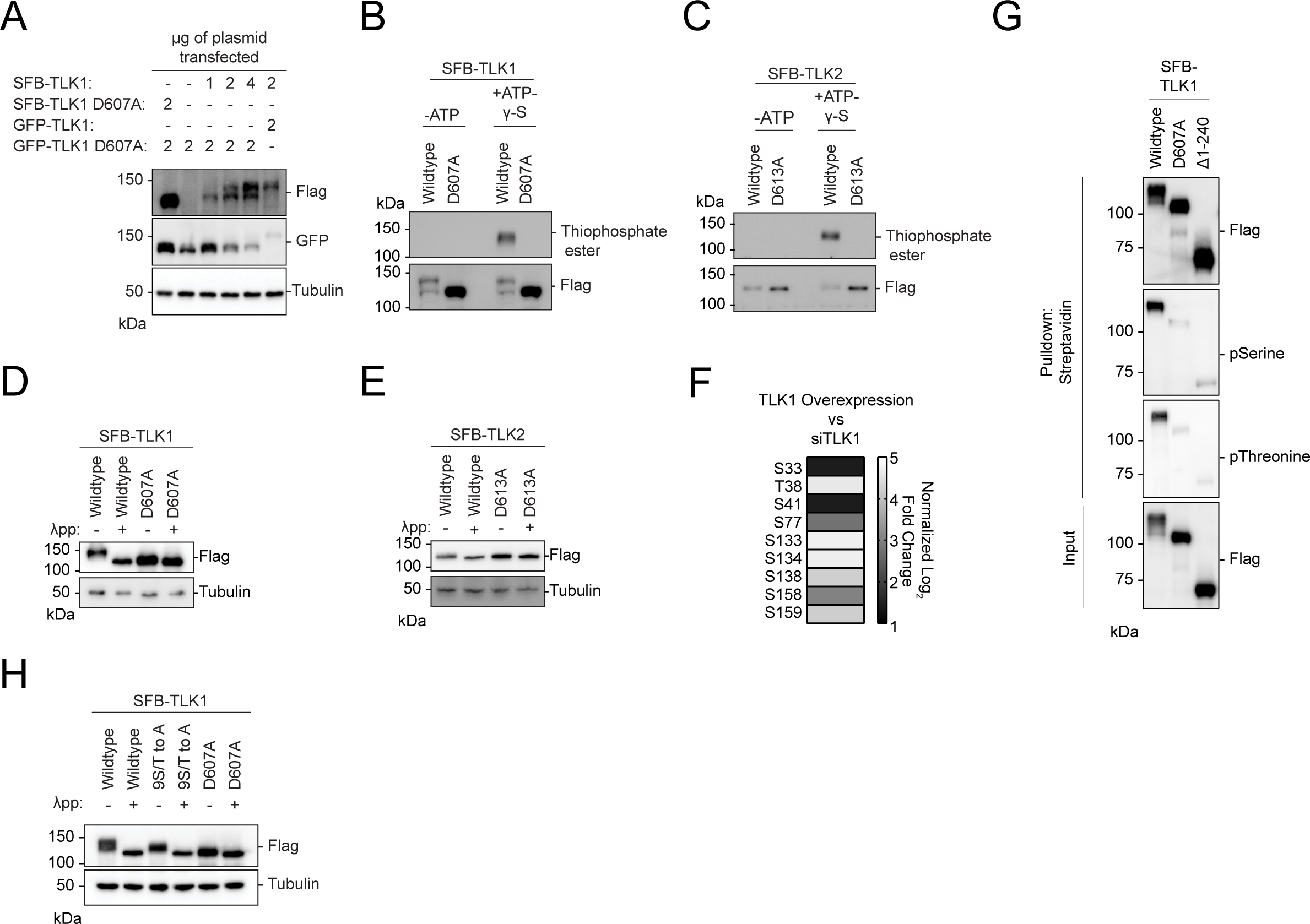
Tousled-like kinases are autophosphorylated at their N-terminus. **(A)** Co-transfection of TLK1 wildtype and catalytic inactive mutant D607A with the indicated amount of plasmids in HEK293T cells followed by Western blotting analysis. **(B-C**) Kinase assays using ATP-γ-S followed by alkylation by p-Nitrobenzyl mesylate to convert phosphorylated residues into thiophosphate esters. Antibody against thiophosphate ester was used to identify phosphorylation events. **(D-E)** Lambda protein phosphatase (λpp) assay was performed using SFB-tagged TLK1 wildtype and D607A mutant or SFB-tagged TLK2 wildtype and D613A mutant purified from HEK293T cell lysate. Differential molecular weights were analyzed by Western blotting analysis. **(F)** TLK1 is autophosphorylated at 9 residues within its N-terminus. Phospho-Tandem-Mass-Tag mass spectrometry was performed on phosphopeptides enriched from HEK293T cells with TLK1 overexpression and knockdown using siRNA. **(G)** TLK1 N-terminus is highly phosphorylated. SFB-tagged TLK1 wildtype, D607A, or Δ1-240 (ΔN-terminus) were purified from HEK293T using streptavidin beads followed by Western blotting analysis using phospho-Serine and phospho-threonine antibodies. **(H)** TLK1 wildtype, N-terminal 9S/T to A, and catalytic inactive mutants were treated with λ-phosphatase. Differential molecular weights were analyzed by Western blot.

Consistently, kinase assays showed that wildtype TLK1 had slower electrophoretic mobility compared to the catalytic inactive TLK1 D607A mutant (**Figure 2B**), which showed no detectable activity (**Figure 2B-C**). To confirm that the increased mobility was due to phosphorylation, wildtype TLK1 was treated with lambda phosphatase. Treated TLK1 showed faster electrophoretic mobility similar to corresponding catalytically inactive mutants (**Figure 2D**), consistent with our previous observations **(Figure 1C-D)**. In contrast, wildtype TLK2 migrated more slowly than the catalytically inactive mutant (**Figure 2C-E**), which may correlate with phosphorylation levels. Using quantitative phospho-enrichment proteomics, we identified nine residues at the TLK1 N-terminus that were phosphorylated (**Figure 2F**). Using phospho-serine and phospho-threonine antibodies, we confirmed that the TLK1 N-terminus is the primary target for auto-phosphorylation (**Figure 2G**). Mutation of the nine serine/threonine residues to alanine (9S/T to A) at the N-terminus increased the electrophoretic mobility (**Figure 2H**) indicating that mutation of 9S/T to A reduces, but does not abolish, autophosphorylation. Together, these data suggest that active TLK1 and TLK2 are autophosphorylated and this is mediated in trans through homo or heterodimerization.

### TLK1 and 2 kinase activity regulates their localization at DNA break sites

LC8, a recently identified DNA repair factor, is recruited to DNA breaks(42). Although LC8 interacts with both TLK1 and TLK2, we did not observe GFP-TLK1 or TLK2 accumulation at DNA damage sites induced by laser microirradiation (**Figure 3A**-**B**). Surprisingly, in both TLK1 and TLK2, the N-terminus, coiled-coil 1 domain deletion, and catalytically inactive mutants robustly accumulated at sites of DNA damage **(Figure 3A)**. In contrast, the N-terminus deletion mutants were primarily cytoplasmic and did not localize to sites of damage (**Figure 3A-B**). The catalytically inactive mutants GFP-TLK1(D607A) and TLK2 (D613A) localized at DNA damage sites within three minutes, with faster kinetics compared to LC8, which localizes to laser-induced damaged DNA at 4-5 mins (**Figure 3C**). Moreover, LC8 did not appear to play a role in regulating TLK1 and TLK2 recruitment to DNA damage as its depletion did not alter their localization kinetics (**Supplementary Figure 3A-H**). Consistently, fractionation experiments showed that catalytically dead mutants of TLK1 and TLK2, but not their wildtype counterparts, were highly enriched in the chromatin fraction (**Figure 3D**). Together, our data suggest that the accumulation of TLK1 and 2 at damaged chromatin is primarily mediated by their N-terminus and inhibited by autophosphorylation that occurs independently of LC8.

**Figure 3.**
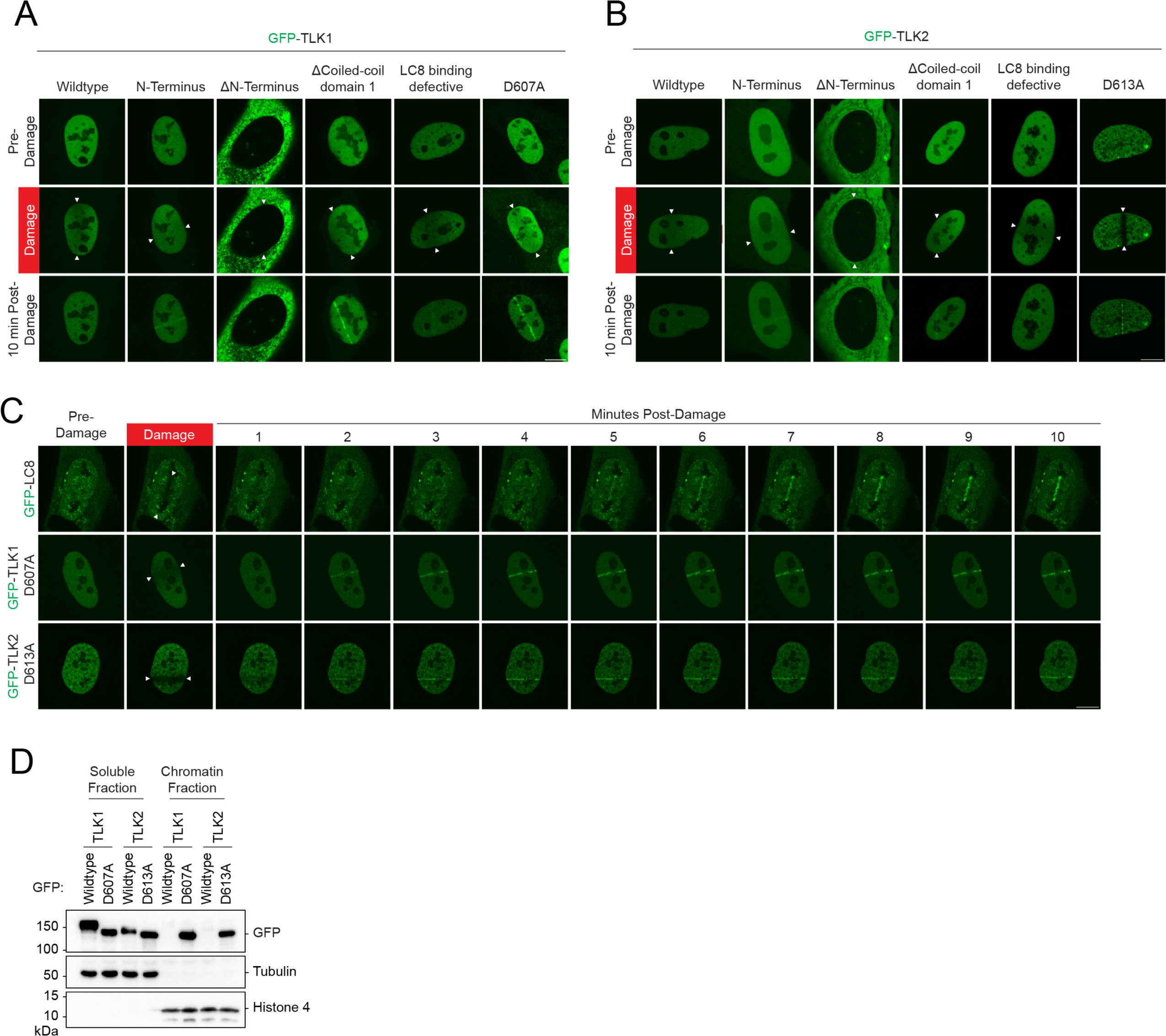
Accumulation of TLK1 and 2 is negatively regulated by their kinase activity and dimerization at damaged chromatin. **(A-B)** TLK1 and TLK2 recruitment to sites of DNA damage is repressed by dimerization and kinase activity. U2OS cells were transfected with indicated GFP-TLK1 or TLK2 wildtype and mutants. Laser-induced micro-irradiation and live-cell imaging experiments were conducted using a confocal microscope, denoted line between the white arrows. Representative images of each experimental group at 10 mins after damage. **(C)** DNA damage kinetics of LC8, TLK1, and TLK2 catalytic inactive mutants using laser-induced micro-irradiation. **(D)** Kinase activity negatively regulates TLK chromatin loading. HEK293T cells were transfected with the indicated GFP-TLK1 wildtype and catalytic inactive mutants. Cells were harvested 24 h after transfection and fractionated to obtain soluble and chromatin fractions.

### N-terminal autophosphorylation inhibits TLK recruitment to DNA damage sites

Given the robust recruitment of the N-terminal fragment and the catalytically inactive mutants of TLK1/2 to laser-induced micro-irradiation, we tested whether the recruitment was dependent on the phosphorylation of the N-terminus. Strikingly, the phosphomimetic mutant (9S/T to D) abolished GFP-TLK1 D607A recruitment to DNA damage sites, whereas the 9S/T to A mutant retained robust recruitment of GFP-TLK1 D607A to DNA damage (**Figure 4A-B**). This suggested that the TLK1 N-terminal auto-phosphorylation sites may be involved in suppressing its retention at DNA damage sites and the regulation is not sequence-specific because of the 9S/T to A D607A mutant can still be recruited to damaged chromatin.

**Figure 4.**
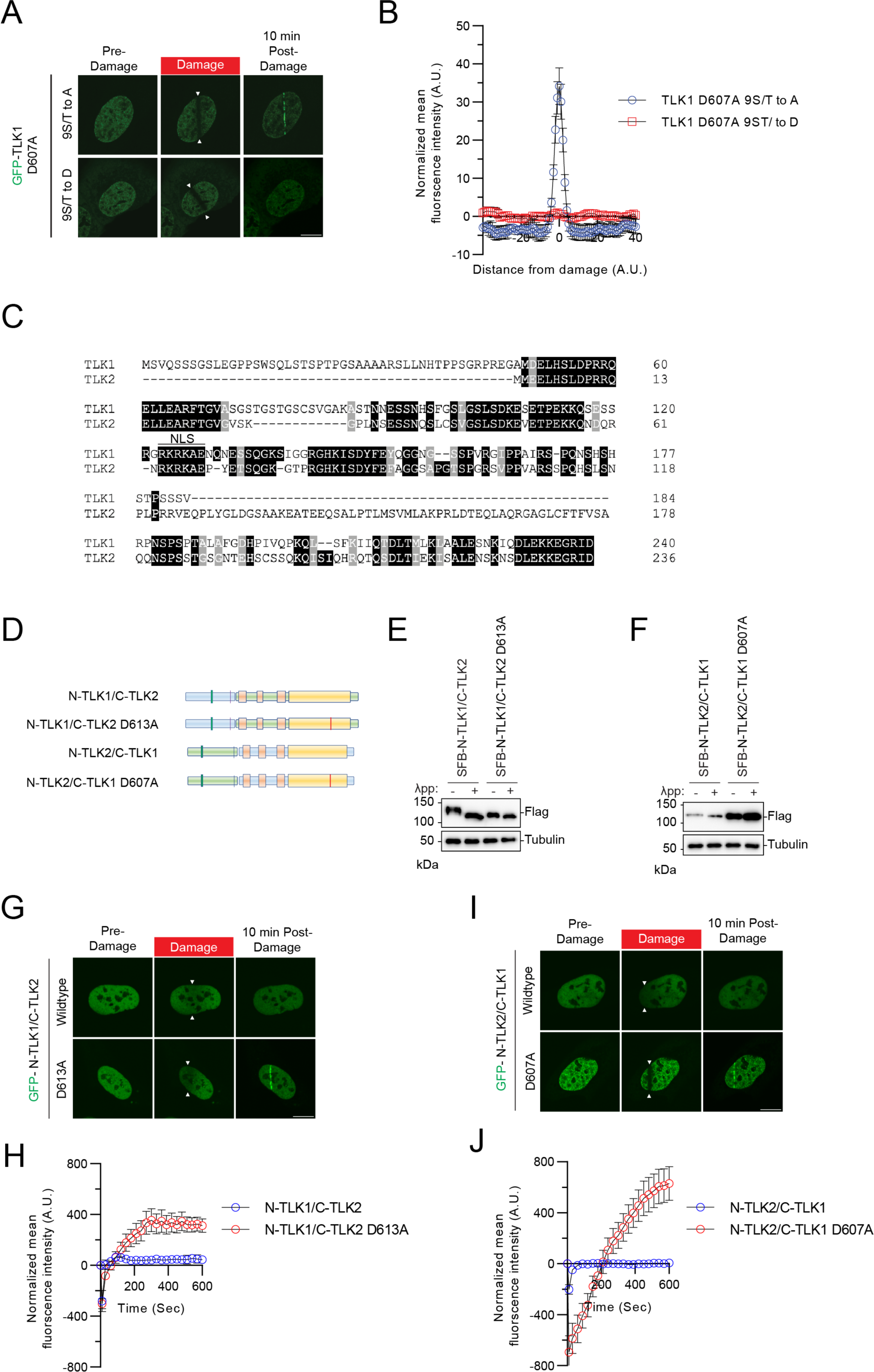
Recruitment of TLK1 and 2 to damaged DNA is N-terminal phosphorylation-dependent. **(A)** TLK1 autophosphorylation suppresses recruitment to damaged chromatin. The TLK1catalytic mutant (D607A) with N-terminus phospho-defective mutant (9S/T to A) or phospho-mimetic mutant (9S/T to D) were subjected to laser-induced micro-irradiation. **(B)** Quantification of the recruitment intensity as shown in A. N=10 **(C)** Sequence alignment of the N-terminus of TLK1 and TLK2. **(D)** Schematic illustration of chimeric proteins generated for TLK1 and TLK2. **(E-F)** Chimeric N-TLK1/C-TLK2, N-TLK1/C-TLK2 D613A, N-TLK2/C-TLK1 and N-TLK2/C-TLK1D607A were treated with λpp. Molecular weights were analyzed by Western blot. **(G)** Representative images of GFP-N-TLK1/C-TLK2 and N-TLK1/C-TLK2 D613A chimeras at 10 mins after laser-induced micro-irradiation. The damaged area was denoted between the arrows. **(H)** Recruitment kinetics quantification of the GFP-chimera proteins after laser-induced micro-irradiation as in (G). n≥10 **(I)** Representative images of GFP-N-TLK2/C-TLK1 and N-TLK2/C-TLK1 D607A chimeras at 10 mins after laser-induced micro-irradiation. The damaged area was denoted between the arrows. **(J)** Recruitment kinetics quantification of the GFP-chimera proteins after laser-induced micro-irradiation as in (I). n≥10

To confirm that the robust mobility shift of the wildtype TLK1 is due to the hyperphosphorylation of its N-terminus, a region where TLK1 and TLK2 share low sequence homology (**Figure 4C**), we performed a domain-swapping experiment by fusing the N-terminus of TLK1 with the TLK2 coiled-coil domains and kinase domain and vice versa (**Figure 4D**). Chimeric protein consisting of TLK1 N-terminus-TLK2 C-terminus (N-TLK1/C-TLK2) showed slower electrophoretic mobility compared to the TLK1 N-terminus-TLK2 C-terminus catalytically inactive (N-TLK1-C-TLK2D613A) chimeric protein, indicating the shift was due to autophosphorylation. Consistent with the lower mobility of wildtype TLK2 compared to TLK1, the N-TLK2/C-TLK1 chimera showed a small mobility difference compared to the N-TLK2/C-TLK1 D607A. Although both TLK1 and TLK2 are auto-phosphorylated at their N-termini, the TLK1 N-terminal hyperphosphorylation results in a more pronounced mobility shift (**Figure 4E-F**). Despite this difference, their N-terminal autophosphorylation consistently regulated their recruitment to DNA damage (**Figure 4G-J**).

To pinpoint the N-terminal domain of TLK1 required for the recruitment to DNA damage sites, we generated systematic serial deletions of TLK1 at the N-terminus, without perturbing the nuclear localization signal, on the wildtype, D607A, and N-terminus (1–240) backbone. Deletion of the first 47 amino acids (Δ47), which is unique to TLK1, or first 116 (Δ116) where TLK1 and TLK2 share some sequence homology (**Figure 5A-B**, **Figure 4C**), showed a modest reduction in TLK1 D607A recruitment to DNA damage. Strikingly, deletion of 133-204 (Δ133-204), where three hyperphosphorylated residues (S133, S134, and S138) reside, completely abolished TLK1 D607A recruitment to DNA damage (**Figure 5C-D**). Consistently, the N-terminus (1–240) Δ133-204 mutant showed a dramatic reduction of recruitment to DNA damage when compared to the wildtype 1-240 fragment (**Supplementary Figure 4A-D**). As expected, the deletion did not alter the DNA damage recruitment of wildtype TLK1 due to the hyperphosphorylation of TLK1 at the basal level (**Supplementary Figure 4E-H**).

**Figure 5.**
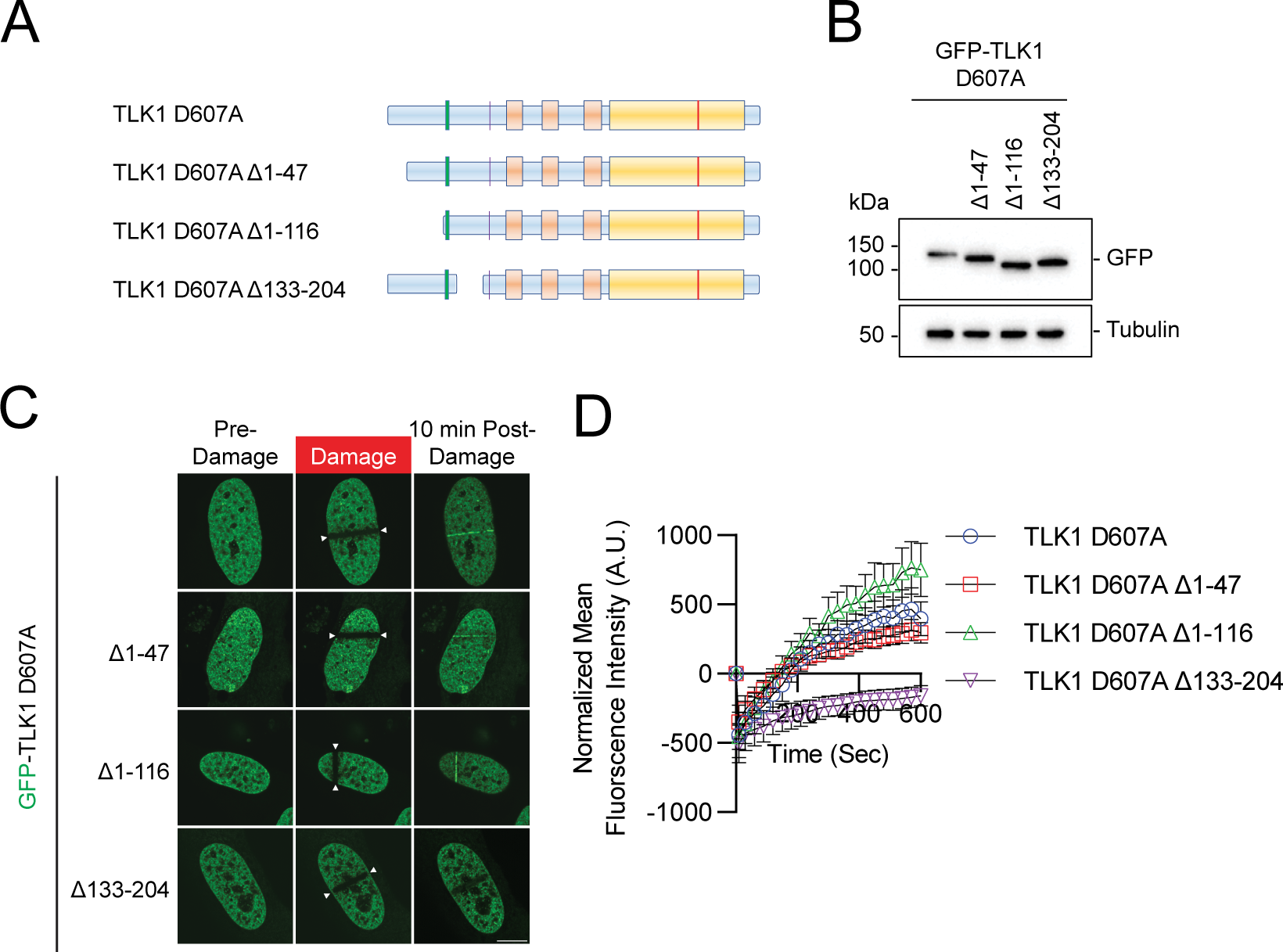
The N-terminus conserved region of TLK1 is required for its laser-induced micro-irradiation. **(A)** Schematic illustrations of TLK1 N-terminal deletion mutants for damaged chromatin recruitment region mapping. **(B)** Western blotting analysis of the mutants as in A and used in C and D. **(C)** Representative images of TLK1 N-terminal deletion mutants 10 mins after laser-induced micro-irradiation. **(D)** Recruitment kinetics of the TLK1 N-terminal deletion mutants as in C. n≥10

### PCNA recruits TLK1 and TLK2 DNA damage sites via protein-protein interaction

To dissect the mechanism by which TLK1 and TLK2 are recruited to damaged chromatin, we used a comparative proteomic approach to identify the upstream mediator for the recruitment of TLK to damaged chromatin. We performed TAP-MS using TLK1 wildtype, D607A mutant, and 1-240 fragments as baits. Consistent with previously reported TLK-PCNA interaction identified by proteomic screens (57), we found that PCNA was enriched in both the TLK1 D607A and 1-240 fragments (**Figure 6A-B**), both of which were recruited to DNA damage induced by laser-induced micro-irradiation (**Figure 3A**). Within the region between aa133-240, sequence analysis showed that both TLK1 and TLK2 contain a non-canonical <u>P</u>CNA interacting protein-<u>b</u>ox (PIP box) ΨxxΣΣ (58, 59) (**Figure 6C**) which is highly evolutionarily conserved (**Figure 6D**). Pulldown assays showed that both the TLK1 and TLK2 catalytically inactive mutants had increased binding affinity to PCNA compared to their wildtype counterpart and mutation of the PIP-box abolished the PCNA binding to the catalytically inactive mutants. TLK2 wildtype showed a low level of PCNA binding, potentially due to its low basal level of hyperphosphorylation at the N-terminus. Furthermore, we were able to detect PCNA with mass spectrometry using a small TLK1 fragment (a.a. 133-208), but not the PIP-box mutant, indicating their specific interaction (**Figure 5**). Notably, the PIP-box mutation abolished the recruitment of TLK1 and TLK2 catalytically inactive mutants to laser-induced micro-irradiation DNA damage (**Figure 6 G-L**), as well as in the small fragment (**Supplementary Figure 5B-C**). Together, our data reveal that N-terminal autophosphorylation of TLK1 and TLK2 inhibited the interaction with PCNA and regulated their recruitment to DNA damage.

**Figure 6.**
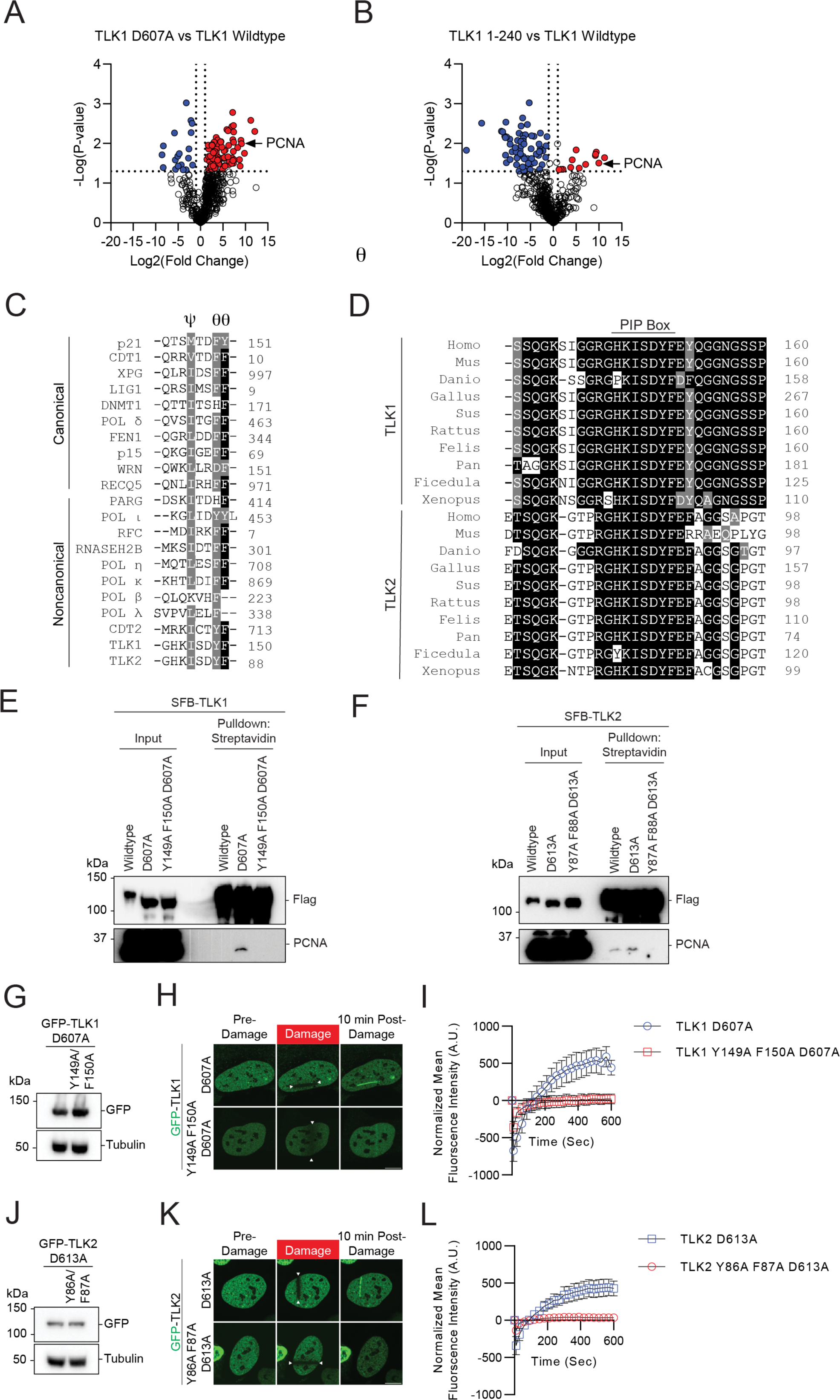
PCNA regulates TLK1 and TLK2 DNA damage recruitment via interaction with their PIP-box. **(A-B)** Volcano plot of proteomic analyses between TLK1 wildtype and catalytic mutant D607A or N-terminal fragment a.a. 1-240. Data was analyzed using Perseus and represents technical duplicate data. **(C)** Sequence alignment of the canonical and non-canonical PIP-box between TLK, TLK2, and other PCNA interacting proteins. PIP-box QxxɸxxΨΨ, where x represents any amino acid, ɸ represents hydrophobic residues, and Ψ represents aromatic residues) or non-canonical sequences (xxxɸxxΨΨ). **(D)** Sequence alignment of the TLK1 and TLK2 PIP-box across species. **(E-F)** TLK1 and TLK2 interact with PCNA. HEK293T cells transfected with SFB-TLK1 wildtype, D607A, or PIP box mutant Y149A F150A D607A. Cells were pulled down by streptavidin beads and analyzed by western blot using indicated antibodies. **(G-I)** TLK1 PIP box is required for DNA damage recruitment. G. GFP-TLK1D607A and D607A PCNA binding defective mutant (Y149A/F150A) protein expression. H. Representative images of indicated mutants at 10 mins after laser-induced micro-irradiation. The irradiated area is induced by the arrows. I. DNA damage recruitment kinetics quantification as in H. N = 10 cells. **(J-L)** TLK2 PIP box is required for DNA damage recruitment. J. GFP-TLK2D613A and D613A PCNA binding defective mutant (Y86A/F87A) protein expression. K. Representative images of indicated mutants at 10 mins after laser-induced micro-irradiation. The irradiated area is induced by the arrows. L. DNA damage recruitment kinetics quantification as in K. n≥10.

## Discussion

Emerging evidence demonstrates that TLK1 and TLK2 are involved in a wide range of cellular processes, including development, DNA replication, cell cycle regulation, and transcription (3, 60). Pathophysiologically, the inactivation of TLK1 and TLK2 leads to DNA damage, innate immune activation, induction of alternative lengthening of telomeres (ALT) pathway, and increased replication stress(1, 6). The functions of TLK1 and TLK2 have been widely investigated through their interactors and substrates, including ASF1A, ASF1B, NEK1, RAD9, DYNLL1/LC8, and RAD54 (4, 8, 28, 30, 39, 54, 60). Yet, the accumulation of TLK1 and TLK2 at DNA damage sites and the mechanisms underlying their chromatin recruitment have never been shown.

In this study, we found that TLK1 and TLK2 accumulate at DNA damage sites, specifically in their inactive state. Their DNA damage recruitment is mediated by interactions with PCNA that are negatively regulated by their dimerization-mediated N-terminal auto-phosphorylation. Notably, TLK1/2 activity is inactivated after multiple types of DNA damage, including IR, UV irradiation, and hydroxyurea treatment (26). We speculate that the activity-dependent interaction mechanism rapidly relocalizes TLK1 and TLK2 to sites of damage and allows them to phosphorylate substrates at the break sites and dynamically dissociate from chromatin. It is also possible that TLK1 and TLK2 have biological functions at damaged chromatin that are independent of their activity. Additionally, TLK1/2 activity is cell cycle-dependent with the highest activity in early S-phase and declining over mid and late S-phase (26), highlighting its role in targeting the ASF1 histone chaperone throughout replication that may not occur on chromatin.

During homologous recombination, PCNA is loaded on the strand invaded D-loop and we speculate that this could result in the recruitment of TLK1 and TLK2 there for the regulation of repair factors at damaged chromatin, including RAD54 and ASF1 (34, 39, 61, 62). In line with a recent study, the hypersensitivity of TLK depletion implicates their potential involvement in homologous recombination repair(1).

Besides the PCNA binding at the TLK1 and TLK2 N-terminus, LC8/DYNLL1 shows constitutive binding through the putative LC8 binding motif on TLK1 (SFKI**<u>IQT</u>**D) and TLK2 (QHRQ**<u>TQS</u>**D). In contrast to the 53BP1-LC8 interaction, there is no apparent regulatory role for LC8 binding to TLK1 or TLK2 in the context of their recruitment to damaged chromatin. Moreover, despite the well-characterized role of LC8 in regulating protein dimerization(56), LC8 is not required for TLK1 and TLK2 dimerization. The functional connection between LC8 and TLKs thus awaits future investigation.

Pathological mutations of TLK1 and TLK2 were previously reported. Mutations of TLK1 at the N-terminus P25L, T38fs, and R142S are associated with intellectual disability and developmental defects (20) and mutations of TLK2, D551G, E475Ter and S617L, are clinically implicated in intellectual disability and developmental defects (9, 63). These TLK2 mutations showed reduced kinase activity, increased DNA damage, chromatin compaction defects, and mis-localization. Interestingly, a TLK2 mutant with coiled-coil domain 1 deletion showed to perinuclear accumulation (54), coinciding with its reduced dimerization and kinase activity and suggesting that these mutants may also have impaired localization to DNA damage site.

TLK1 and TLK2 appear to constitutively form higher-order oligomers (54). However, it is unclear how they stoichiometrically dimerize or oligomerize in physiological and pathological conditions. As we observed that dimerization is essential for their autophosphorylation, we speculate that at least the initial activation and phosphorylation occurs in trans and the constitutive autophosphorylation is maintained by both in trans and in cis activity. There is no report showing that TLK1 and 2 function as monomers in cells and it is likely that their reduced activity after DNA damage and mid to late S phase is regulated by CHK1 activity rather than dimer dissociation (26).

There are ongoing efforts to develop TLK1 and TLK2 inhibitors for targeting cancer cells with their promising roles in maintaining genome stability (64–66). Emerging evidence demonstrated that TLK1 and TLK2 are involved in cancer progression and therapeutic resistance (67–70) and it has been shown that inhibition of TLK potentiates cell-killing in different types of cancers (1, 70–74), making TLK 1 and 2 attractive targets for therapeutic strategy development.

Overall, our study provides strong evidence that TLK1 and 2 accumulate at DNA damage sites via autophosphorylation-mediated PCNA interaction. This biological regulation can potentially be used to manipulate TLK function to sensitize cells to radiation and chemotherapeutic agents by targeting the DNA damage response pathway.

## Data availability

The proteomic data for this study have been uploaded to ProteomeXchange, project accession numbers: PXD047964, PXD047966, PXD047996 and PXD048215.

## Acknowledgments

We thank Dr. Travis Stracker for insightful discussion and critical reading of the manuscript. We would also like to thank Kyle Tengler and Dr. Tram Nguyen for technical supports on reagent verification.

## Author contributions

J.L. and G.G. conceived and designed the study. K.L.W., N.K., G.G. and J.L. performed the experiments and analyzed the data. J.L. and K.L.W. wrote the manuscript, G.G. and K.R. reviewed and edited the paper with input from other authors.

## References

1. Lee, S.B., Segura-Bayona, S., Villamor-Payà, M., Saredi, G., Todd, M.A.M., Attolini, C.S.O., Chang, T.Y., Stracker, T.H. and Groth, A. (2018) Tousled-like kinases stabilize replication forks and show synthetic lethality with checkpoint and PARP inhibitors. Science Advances, 4.

2. Li, Y., DeFatta, R., Anthony, C., Sunavala, G. and De Benedetti, A. (2001) A translationally regulated Tousled kinase phosphorylates histone H3 and confers radioresistance when overexpressed. Oncogene, 20, 726–738.

3. Segura-Bayona, S. and Stracker, T.H. (2019) The Tousled-like kinases regulate genome and epigenome stability: implications in development and disease. Cell Mol Life Sci, 76, 3827–3841.

4. Canfield, C., Rains, J. and De Benedetti, A. (2009) TLK1B promotes repair of DSBs via its interaction with Rad9 and Asf1. BMC Molecular Biology, 10, 110.

5. Sunavala-Dossabhoy, G., Balakrishnan, K., Sen, S., Nuthalapaty, S. and De Benedetti, A. (2005) The radioresistance kinase TLK1B protects the cells by promoting repair of double strand breaks. BMC Molecular Biology, 6.

6. Segura-Bayona, S., Villamor-Payà, M., Attolini, C.S.O., Koenig, L.M., Sanchiz-Calvo, M., Boulton, S.J. and Stracker, T.H. (2020) Tousled-Like Kinases Suppress Innate Immune Signaling Triggered by Alternative Lengthening of Telomeres. Cell Reports, 32.

7. Segura-Bayona, S., Knobel, P.A., Gonzalez-Buron, H., Youssef, S.A., Peña-Blanco, A., Coyaud, E., Lopez-Rovira, T., Rein, K., Palenzuela, L., Colombelli, J., et al. (2017) Differential requirements for Tousled-like kinases 1 and 2 in mammalian development. Cell Death and Differentiation, 24, 1872–1885.

8. Silljé, H.H.W. and Nigg, E.A. (2001) Identification of human Asf1 chromatin assembly factors as substrates of Tousled-like kinases. Current Biology, 11, 1068–1073.

9. Pavinato, L., Villamor-Payà, M., Sanchiz-Calvo, M., Andreoli, C., Gay, M., Vilaseca, M., Arauz-Garofalo, G., Ciolfi, A., Bruselles, A., Pippucci, T., et al. (2022) Functional analysis of TLK2 variants and their proximal interactomes implicates impaired kinase activity and chromatin maintenance defects in their pathogenesis. J Med Genet, 59, 170–179.

10. Han, Z., Saam, J.R., Adams, H.P., Mango, S.E. and Schumacher, J.M. (2003) The C. elegans Tousled-like Kinase (TLK-1) Has an Essential Role in Transcription. Current Biology, 13, 1921–1929.

11. Lee, J., Kim, M.S., Park, S.H. and Jang, Y.K. (2018) Tousled-like kinase 1 is a negative regulator of core transcription factors in murine embryonic stem cells. Sci Rep, 8, 334.

12. Sillje, H.H.W., Takahashi, K., Van Houwe, G. and Nigg, E.A. (1999) Mammalian homologues of the plant Tousled gene code for cell-cycle-regulated kinases with maximal activities linked to ongoing DNA replication. The EMBO Journal, 18, 5691–5702.

13. Kim, J.-A., Anurag, M., Veeraraghavan, J., Schiff, R., Li, K. and Wang, X. (2016) Amplification of TLK2 Induces Genomic Instability via Impairing the G2/M Checkpoint. Molecular Cancer Research, 14, 920–927.

14. Han, Z., Riefler, G.M., Saam, J.R., Mango, S.E. and Schumacher, J.M. (2005) The C. elegans Tousled-like Kinase Contributes to Chromosome Segregation as a Substrate and Regulator of the Aurora B Kinase. Current Biology, 15, 894–904.

15. Bruinsma, W., van den Berg, J., Aprelia, M. and Medema, R.H. (2016) Tousled-like kinase 2 regulates recovery from a DNA damage-induced G2 arrest. EMBO Rep, 17, 659–670.

16. Awate, S. and De Benedetti, A. (2016) TLK1B mediated phosphorylation of Rad9 regulates its nuclear/cytoplasmic localization and cell cycle checkpoint. BMC Mol Biol, 17, 3.

17. Roe, J.L., Rivin, C.J., Sessions, R.A., Feldmann, K.A. and Zambryski, P.C. (1993) The Tousled gene in A. thaliana encodes a protein kinase homolog that is required for leaf and flower development. Cell, 75, 939–950.

18. Yeh, T.-H., Huang, S.-Y., Lan, W.-Y., Liaw, G.-J. and Yu, J.-Y. (2015) Modulation of Cell Morphogenesis by Tousled-Like Kinase in the Drosophila Follicle Cell. Dev Dyn, 10.1002/dvdy.

19. Carrera, P., Moshkin, Y.M., Grönke, S., Silljé, H.H.W., Nigg, E.A., Jäckle, H. and Karch, F. (2003) Tousled-like kinase functions with the chromatin assembly pathway regulating nuclear divisions. Genes Dev, 17, 2578–2590.

20. Villamor-Payà, M., Sanchiz-Calvo, M., Smak, J., Pais, L., Sud, M., Shankavaram, U., Lovgren, A.K., Austin-Tse, C., Ganesh, V.S., Gay, M., et al. (2023) Identification of a de novo mutation in TLK1 associated with a neurodevelopmental disorder and immunodeficiency. medRxiv, 10.1101/2023.08.22.23294267, 24 August 2023, pre-print: not peer-reviewed.

21. Reijnders, M.R.F., Janowski, R., Alvi, M., Self, J.E., van Essen, T.J., Vreeburg, M., Rouhl, R.P.W., Stevens, S.J.C., Stegmann, A.P.A., Schieving, J., et al. (2018) PURA syndrome: clinical delineation and genotype-phenotype study in 32 individuals with review of published literature. J Med Genet, 55, 104–113.

22. Töpf, A., Oktay, Y., Balaraju, S., Yilmaz, E., Sonmezler, E., Yis, U., Laurie, S., Thompson, R., Roos, A., MacArthur, D.G., et al. (2020) Severe neurodevelopmental disease caused by a homozygous TLK2 variant. Eur J Hum Genet, 28, 383–387.

23. Woods, E., Spiller, M. and Balasubramanian, M. (2022) Report of two children with global developmental delay in association with de novo TLK2 variant and literature review. Am J Med Genet A, 188, 931–940.

24. Lelieveld, S.H., Reijnders, M.R.F., Pfundt, R., Yntema, H.G., Kamsteeg, E.-J., de Vries, P., de Vries, B.B.A., Willemsen, M.H., Kleefstra, T., Löhner, K., et al. (2016) Meta-analysis of 2, 104 trios provides support for 10 new genes for intellectual disability. Nat Neurosci, 19, 1194–1196.

25. Ehsan, H., Reichheld, J.-P., Durfee, T. and Roe, J.L. (2004) TOUSLED Kinase Activity Oscillates during the Cell Cycle and Interacts with Chromatin Regulators. Plant Physiol, 134, 1488–1499.

26. Groth, A., Lukas, J., Nigg, E.A., Silljé, H.H.W., Wernstedt, C., Bartek, J. and Hansen, K. (2003) Human Tousled like kinases are targeted by an ATM-and Chk1-dependent DNA damage checkpoint. EMBO J, 22, 1676–1687.

27. Krause, D.R., Jonnalagadda, J.C., Gatei, M.H., Sillje, H.H.W., Zhou, B.-B., Nigg, E.A. and Khanna, K. (2003) Suppression of Tousled-like kinase activity after DNA damage or replication block requires ATM, NBS1 and Chk1. Oncogene, 22, 5927–5937.

28. Simon, B., Lou, H.J., Huet-Calderwood, C., Shi, G., Boggon, T.J., Turk, B.E. and Calderwood, D.A. (2022) Tousled-like kinase 2 targets ASF1 histone chaperones through client mimicry. Nat Commun, 13, 749.

29. Pilyugin, M., Demmers, J., Verrijzer, C.P., Karch, F. and Moshkin, Y.M. (2009) Phosphorylation-Mediated Control of Histone Chaperone ASF1 Levels by Tousled-Like Kinases. PLoS One, 4, e8328.

30. Klimovskaia, I.M., Young, C., Strømme, C.B., Menard, P., Jasencakova, Z., Mejlvang, J., Ask, K., Ploug, M., Nielsen, M.L., Jensen, O.N., et al. (2014) Tousled-like kinases phosphorylate Asf1 to promote histone supply during DNA replication. Nat Commun, 5, 3394.

31. Huang, T.H., Fowler, F., Chen, C.C., Shen, Z.J., Sleckman, B. and Tyler, J.K. (2018) The Histone Chaperones ASF1 and CAF-1 Promote MMS22L-TONSL-Mediated Rad51 Loading onto ssDNA during Homologous Recombination in Human Cells. Molecular Cell, 69, 879–892.e5.

32. Huang, T.H., Shen, Z.J., Sleckman, B.P. and Tyler, J.K. (2018) The histone chaperone ASF1 regulates the activation of ATM and DNA-PKcs in response to DNA double-strand breaks. Cell Cycle, 17, 1413–1424.

33. Kim, J.A. and Haber, J.E. (2009) Chromatin assembly factors Asf1 and CAF-1 have overlapping roles in deactivating the DNA damage checkpoint when DNA repair is complete. Proc Natl Acad Sci U S A, 106, 1151–1156.

34. Lee, K.Y., Im, J.-S., Shibata, E. and Dutta, A. (2017) ASF1a Promotes Non-homologous End Joining Repair by Facilitating Phosphorylation of MDC1 by ATM at Double-Strand Breaks. Molecular Cell, 68, 61–75.e5.

35. Mello, J.A., Silljé, H.H.W., Roche, D.M.J., Kirschner, D.B., Nigg, E.A. and Almouzni, G. (2002) Human Asf1 and CAF-1 interact and synergize in a repair-coupled nucleosome assembly pathway. EMBO Rep, 3, 329–334.

36. Singh, V., Connelly, Z.M., Shen, X. and De Benedetti, A. (2017) Identification of the proteome complement of humanTLK1 reveals it binds and phosphorylates NEK1 regulating its activity. Cell Cycle, 16, 915–926.

37. Sunavala-Dossabhoy, G. and De Benedetti, A. (2009) Tousled homolog, TLK1, binds and phosphorylates Rad9; TLK1 acts as a molecular chaperone in DNA repair. DNA Repair, 8, 87–102.

38. Kelly, R. and Davey, S.K. (2013) Tousled-Like Kinase-Dependent Phosphorylation of Rad9 Plays a Role in Cell Cycle Progression and G2/M Checkpoint Exit. PLoS ONE, 8, e85859.

39. Ghosh, I., Kwon, Y., Shabestari, A.B., Chikhale, R., Chen, J., Wiese, C., Sung, P. and De Benedetti, A. (2023) TLK1-mediated RAD54 phosphorylation spatio-temporally regulates Homologous Recombination Repair. Nucleic Acids Res, 51, 8643–8662.

40. Khalil, M.I., Madere, C., Ghosh, I., Adam, R.M. and De Benedetti, A. (2021) Interaction of TLK1 and AKTIP as a Potential Regulator of AKT Activation in Castration-Resistant Prostate Cancer Progression. Pathophysiology, 28, 339–354.

41. Wessel, S.R., Mohni, K.N., Luzwick, J.W., Dungrawala, H. and Cortez, D. (2019) Functional Analysis of the Replication Fork Proteome Identifies BET Proteins as PCNA Regulators. Cell Rep, 28, 3497–3509.e4.

42. West, K.L., Kelliher, J.L., Xu, Z., An, L., Reed, M.R., Eoff, R.L., Wang, J., Huen, M.S.Y. and Leung, J.W.C. (2019) LC8/DYNLL1 is a 53BP1 effector and regulates checkpoint activation. Nucleic Acids Res, 47, 6236–6249.

43. Leung, J.W.C., Makharashvili, N., Agarwal, P., Chiu, L.Y., Pourpre, R., Cammarata, M.B., Cannon, J.R., Sherker, A., Durocher, D., Brodbelt, J.S., et al. (2017) ZMYM3 regulates BRCA1 localization at damaged chromatin to promote DNA repair. Genes Dev, 31, 260–274.

44. Ahn, J.H., Davis, E.S., Daugird, T.A., Zhao, S., Quiroga, I.Y., Uryu, H., Li, J., Storey, A.J., Tsai, Y.-H., Keeley, D.P., et al. (2021) Phase separation drives aberrant chromatin looping and cancer development. Nature, 595.

45. Dong, C., West, K.L., Tan, X.Y., Li, J., Ishibashi, T., Yu, C.H., Sy, S.M.H., Leung, J.W.C. and Huen, M.S.Y. (2020) Screen identifies DYRK1B network as mediator of transcription repression on damaged chromatin. Proc Natl Acad Sci U S A, 117, 17019–17030.

46. Zybailov, B.L., Glazko, G.V., Rahmatallah, Y., Andreyev, D.S., McElroy, T., Karaduta, O., Byrum, S.D., Orr, L., Tackett, A.J., Mackintosh, S.G., et al. (2019) Metaproteomics reveals potential mechanisms by which dietary resistant starch supplementation attenuates chronic kidney disease progression in rats. PLoS ONE, 14, e0199274.

47. Cox, J. and Mann, M. (2008) MaxQuant enables high peptide identification rates, individualized p.p.b.-range mass accuracies and proteome-wide protein quantification. Nat Biotechnol, 26, 1367–1372.

48. Storey, A.J., Naceanceno, K.S., Lan, R.S., Washam, C.L., Orr, L.M., Mackintosh, S.G., Tackett, A.J., Edmondson, R.D., Wang, Z., Li, H., et al. (2020) ProteoViz: a tool for the analysis and interactive visualization of phosphoproteomics data. Mol. Omics, 16, 316–326.

49. Ritchie, M.E., Phipson, B., Wu, D., Hu, Y., Law, C.W., Shi, W. and Smyth, G.K. (2015) limma powers differential expression analyses for RNA-sequencing and microarray studies. Nucleic Acids Res, 43, e47–e47.

50. Chen, Y., Han, H., Seo, G., Vargas, R.E., Yang, B., Chuc, K., Zhao, H. and Wang, W. (2020) Systematic analysis of the Hippo pathway organization and oncogenic alteration in evolution. Sci Rep, 10, 3173.

51. He, Y.J., Meghani, K., Caron, M.C., Yang, C., Ronato, D.A., Bian, J., Sharma, A., Moore, J., Niraj, J., Detappe, A., et al. (2018) DYNLL1 binds to MRE11 to limit DNA end resection in BRCA1-deficient cells. Nature, 563, 522–526.

52. Becker, J.R., Cuella-Martin, R., Barazas, M., Liu, R., Oliveira, C., Oliver, A.W., Bilham, K., Holt, A.B., Blackford, A.N., Heierhorst, J., et al. (2018) The ASCIZ-DYNLL1 axis promotes 53BP1-dependent non-homologous end joining and PARP inhibitor sensitivity. Nat Commun, 9.

53. Swift, M.L., Zhou, R., Syed, A., Moreau, L.A., Tomasik, B., Tainer, J.A., Konstantinopoulos, P.A., D’Andrea, A.D., He, Y.J. and Chowdhury, D. (2023) Dynamics of the DYNLL1–MRE11 complex regulate DNA end resection and recruitment of Shieldin to DSBs. Nat Struct Mol Biol, 30, 1456–1467.

54. Mortuza, G.B., Hermida, D., Pedersen, A.-K., Segura-Bayona, S., López-Méndez, B., Redondo, P., Rüther, P., Pozdnyakova, I., Garrote, A.M., Muñoz, I.G., et al. (2018) Molecular basis of Tousled-Like Kinase 2 activation. Nat Commun, 9, 2535.

55. Roe, J.L., Durfee, T., Zupan, J.R., Repetti, P.P., McLean, B.G. and Zambryski, P.C. (1997) TOUSLED is a nuclear serine/threonine protein kinase that requires a coiled-coil region for oligomerization and catalytic activity. J Biol Chem, 272, 5838–5845.

56. Barbar, E. (2008) Dynein light chain LC8 is a dimerization hub essential in diverse protein networks. Biochemistry, 47, 503–508.

57. Srivastava, M., Chen, Z., Zhang, H., Tang, M., Wang, C., Jung, S.Y. and Chen, J. (2018) Replisome Dynamics and Their Functional Relevance upon DNA Damage through the PCNA Interactome. Cell Reports, 25, 3869–3883.e4.

58. Warbrick, E. (1998) PCNA binding proteins in Drosophila melanogaster: the analysis of a conserved PCNA binding domain. Nucleic Acids Res, 26, 3925–3932.

59. Horsfall, A.J., Vandborg, B.A., Kowalczyk, W., Chav, T., Scanlon, D.B., Abell, A.D. and Bruning, J.B. (2021) Unlocking the PIP-box: A peptide library reveals interactions that drive high-affinity binding to human PCNA. J Biol Chem, 296, 100773.

60. Ghosh, I. and De Benedetti, A. (2023) Untousling the Role of Tousled-like Kinase 1 in DNA Damage Repair. Int J Mol Sci, 24, 13369.

61. Tang, M., Chen, Z., Wang, C., Feng, X., Lee, N., Huang, M., Zhang, H., Li, S., Xiong, Y. and Chen, J. (2022) Histone chaperone ASF1 acts with RIF1 to promote DNA end joining in BRCA1-deficient cells. J Biol Chem, 298, 101979.

62. Feng, S., Ma, S., Li, K., Gao, S., Ning, S., Shang, J., Guo, R., Chen, Y., Blumenfeld, B., Simon, I., et al. (2022) RIF1-ASF1-mediated high-order chromatin structure safeguards genome integrity. Nat Commun, 13, 957.

63. Satterstrom, F.K., Kosmicki, J.A., Wang, J., Breen, M.S., De Rubeis, S., An, J.-Y., Peng, M., Collins, R., Grove, J., Klei, L., et al. (2020) Large-Scale Exome Sequencing Study Implicates Both Developmental and Functional Changes in the Neurobiology of Autism. Cell, 180, 568–584.e23.

64. Bhoir, S. and De Benedetti, A. (2023) Targeting Prostate Cancer, the ‘Tousled Way’. Int J Mol Sci, 24, 11100.

65. Johnson, D., Hussain, J., Bhoir, S., Chandrasekaran, V., Sahrawat, P., Hans, T., Khalil, M.I., De Benedetti, A., Thiruvenkatam, V. and Kirubakaran, S. (2023) Synthesis, kinetics and cellular studies of new phenothiazine analogs as potent human-TLK inhibitors. Org Biomol Chem, 21, 1980–1991.

66. Lee, S.-B., Chang, T.-Y., Lee, N.-Z., Yu, Z.-Y., Liu, C.-Y. and Lee, H.-Y. (2022) Design, synthesis and biological evaluation of bisindole derivatives as anticancer agents against Tousled-like kinases. Eur J Med Chem, 227, 113904.

67. Khalil, M.I. and De Benedetti, A. (2022) Tousled-like kinase 1: a novel factor with multifaceted role in mCRPC progression and development of therapy resistance. Cancer Drug Resist, 5, 93–101.

68. Khalil, M.I. and De Benedetti, A. (2022) The TLK1-MK5 Axis Regulates Motility, Invasion, and Metastasis of Prostate Cancer Cells. Cancers (Basel), 14, 5728.

69. Zhang, Z. and Liu, S. (2023) The interaction between ASF1B and TLK1 promotes the malignant progression of low-grade glioma. Ann Med, 55, 1111–1122.

70. Kim, J.-A., Tan, Y., Wang, X., Cao, X., Veeraraghavan, J., Liang, Y., Edwards, D.P., Huang, S., Pan, X., Li, K., et al. (2016) Comprehensive functional analysis of the tousled-like kinase 2 frequently amplified in aggressive luminal breast cancers. Nat Commun, 7.

71. Ibrahim, K., Abdul Murad, N.A., Harun, R. and Jamal, R. (2020) Knockdown of Tousled-like kinase 1 inhibits survival of glioblastoma multiforme cells. Int J Mol Med, 46, 685–699.

72. Ronald, S., Awate, S., Rath, A., Carroll, J., Galiano, F., Dwyer, D., Kleiner-Hancock, H., Mathis, J.M., Vigod, S. and De Benedetti, A. (2013) Phenothiazine Inhibitors of TLKs Affect Double-Strand Break Repair and DNA Damage Response Recovery and Potentiate Tumor Killing with Radiomimetic Therapy. Genes Cancer, 4, 39– 53.

73. Takayama, Y., Kokuryo, T., Yokoyama, Y., Ito, S., Nagino, M., Hamaguchi, M. and Senga, T. (2010) Silencing of Tousled-like kinase 1 sensitizes cholangiocarcinoma cells to cisplatin-induced apoptosis. Cancer Lett, 296, 27– 34.

74. Ronald, S., Sunavala-Dossabhoy, G., Adams, L., Williams, B. and De Benedetti, A. (2011) The expression of tousled kinases in CaP cell lines and its relation to radiation response and DSB repair. The Prostate, 71, 1367–1373.

