## Supplemental figure 1.5 for "Autophosphorylation of the Tousled-like kinases TLK1 and TLK2 regulates recruitment to damaged chromatin via PCNA interaction"

### Figure S1

A

| Identified Protein | Spectral Counts |
|---|---|
| TLK1 (Bait) | 645 |
| TLK2 | 194 |
| PGAM5 | 26 |
| HSPA1B | 22 |
| HSPA8 | 20 |
| HSPA5 | 16 |
| LC8 | 15 |
| HNRNPM | 14 |
| RPL11 | 14 |
| ACACA | 12 |

| Identified Protein | Spectral Counts |
|---|---|
| TLK2 (Bait) | 1296 |
| TLK1 | 251 |
| TUBB | 114 |
| HSPA8 | 91 |
| HSPA1B | 48 |
| HSPA5 | 43 |
| LC8 | 41 |
| ACACA | 30 |
| MCCC2 | 29 |
| HNRNPM | 23 |

B

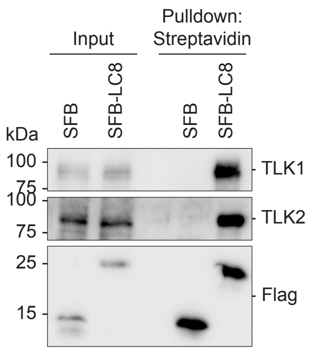


C

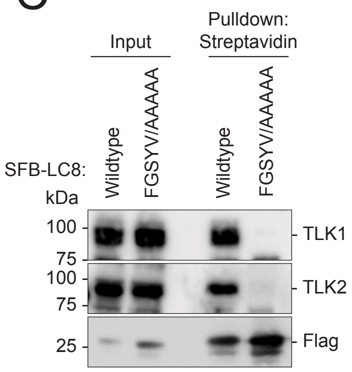


D

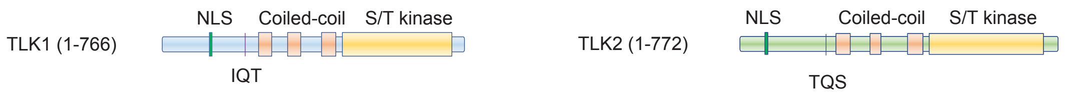


E

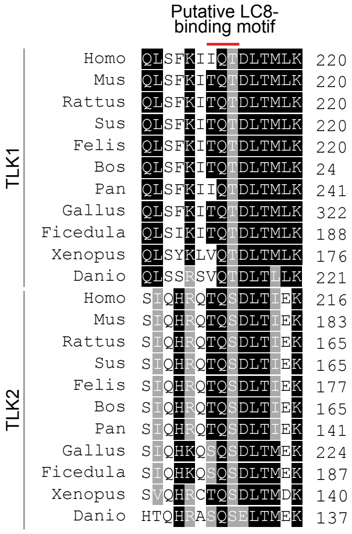


F

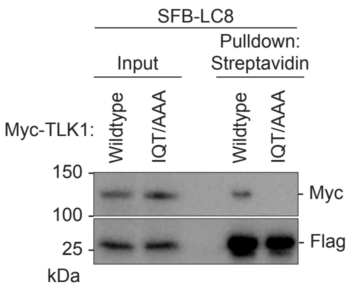


G

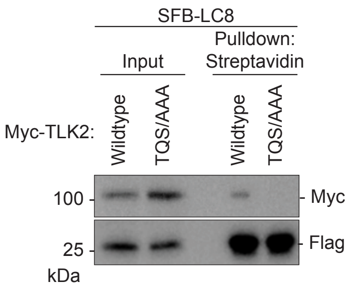


H

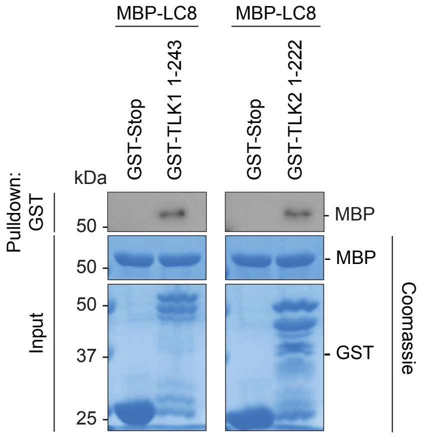

### Figure S2

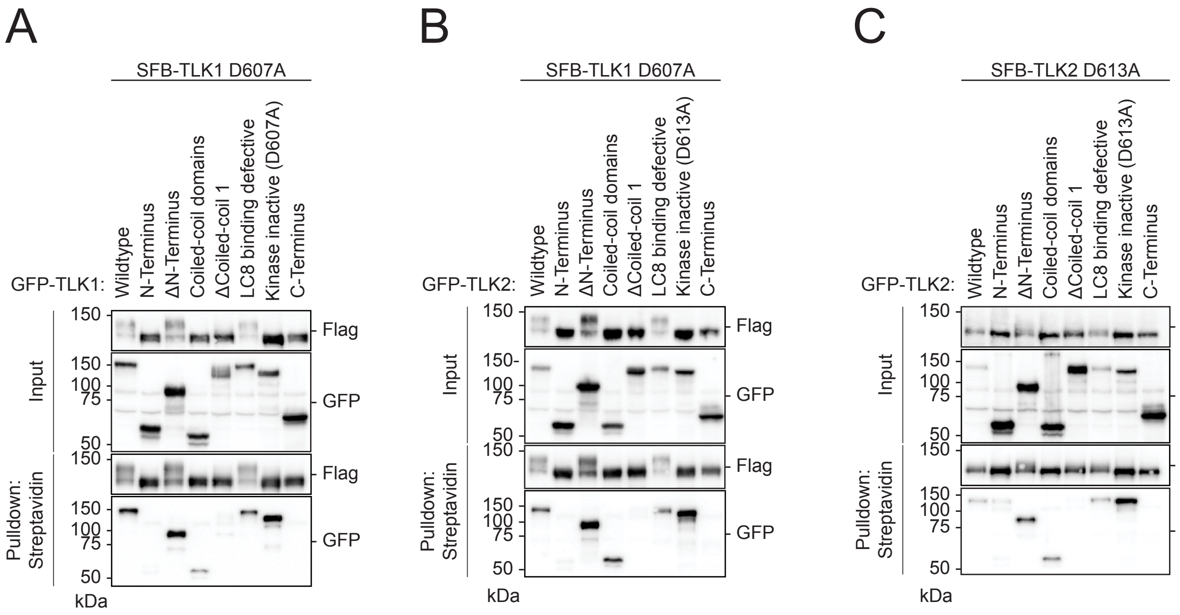
A
SFB-TLK1 D607A
GFP-TLK1:
Wildtype
N-Terminus
ΔN-Terminus
Coiled-coil domains
ΔCoiled-coil 1
LC8 binding defective
Kinase inactive (D607A)
C-Terminus
Input
Pulldown: Streptavidin
Flag
GFP
150
100
75
50
kDa
B
SFB-TLK1 D607A
GFP-TLK2:
Kinase inactive (D613A)
C
SFB-TLK2 D613A
GFP-TLK2:

Figure S3

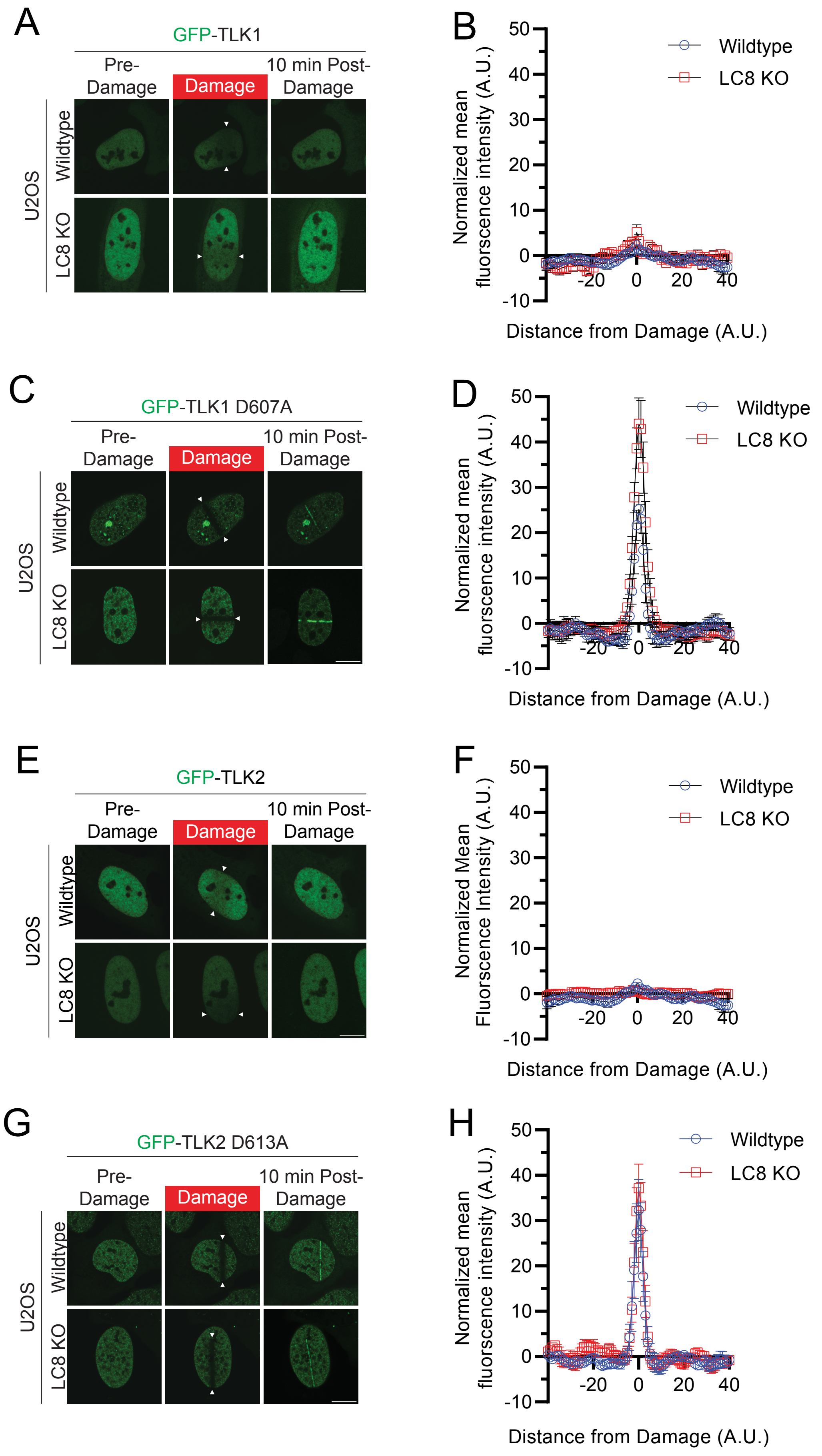
A
GFP-TLK1
Pre-Damage
Damage
10 min Post-Damage
U2OS
Wildtype
LC8 KO
B
Normalized mean fluorescence intensity (A.U.)
Distance from Damage (A.U.)
Wildtype
LC8 KO
50
40
30
20
10
0
-10
-20
0
20
40
C
GFP-TLK1 D607A
Pre-Damage
Damage
10 min Post-Damage
U2OS
Wildtype
LC8 KO
D
Normalized mean fluorescence intensity (A.U.)
Distance from Damage (A.U.)
Wildtype
LC8 KO
E
GFP-TLK2
Pre-Damage
Damage
10 min Post-Damage
U2OS
Wildtype
LC8 KO
F
Normalized Mean Fluorescence Intensity (A.U.)
Distance from Damage (A.U.)
Wildtype
LC8 KO
G
GFP-TLK2 D613A
Pre-Damage
Damage
10 min Post-Damage
U2OS
Wildtype
LC8 KO
H
Normalized mean fluorescence intensity (A.U.)
Distance from Damage (A.U.)
Wildtype
LC8 KO

### Figure S4

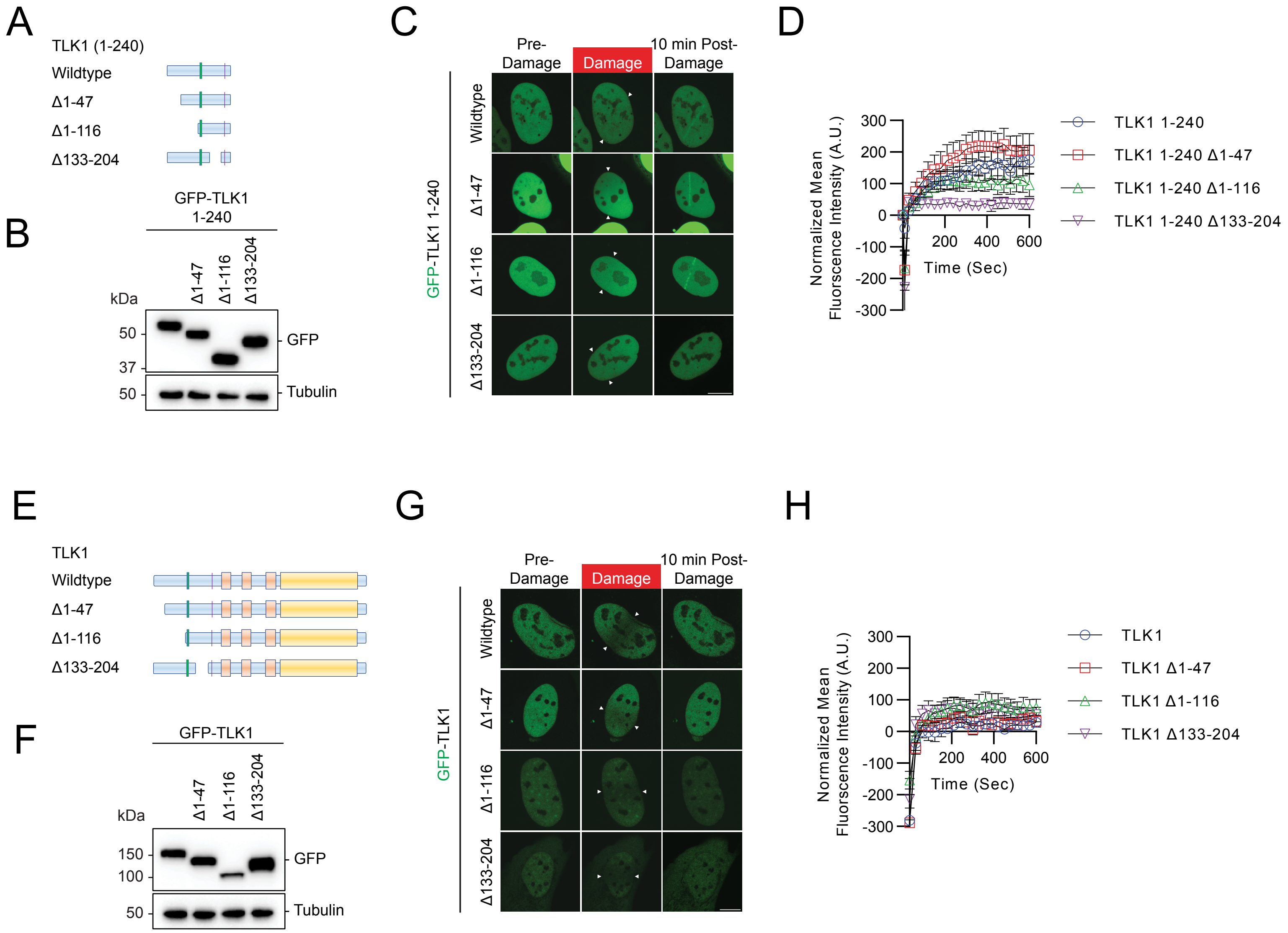

A
TLK1 (1-240)
Wildtype
Δ1-47
Δ1-116
Δ133-204
B
GFP-TLK1
1-240
Δ1-47
Δ1-116
Δ133-204
kDa
50
37
50
GFP
Tubulin
C
Pre-
Damage
Damage
10 min Post-
Damage
GFP-TLK1 1-240
Wildtype
Δ1-47
Δ1-116
Δ133-204
D
Normalized Mean
Fluorscence Intensity (A.U.)
300
200
100
0
-100
-200
-300
200
400
600
Time (Sec)
TLK1 1-240
TLK1 1-240 Δ1-47
TLK1 1-240 Δ1-116
TLK1 1-240 Δ133-204
E
TLK1
Wildtype
Δ1-47
Δ1-116
Δ133-204
F
GFP-TLK1
Δ1-47
Δ1-116
Δ133-204
kDa
150
100
50
GFP
Tubulin
G
Pre-
Damage
Damage
10 min Post-
Damage
GFP-TLK1
Wildtype
Δ1-47
Δ1-116
Δ133-204
H
Normalized Mean
Fluorscence Intensity (A.U.)
300
200
100
0
-100
-200
-300
200
400
600
Time (Sec)
TLK1
TLK1 Δ1-47
TLK1 Δ1-116
TLK1 Δ133-204

### Figure S5

A

| | Spectral Counts | |
|---|---|---|
| Identified Protein | TLK1 133 - 208 | TLK1 133-208 Y149A F150A |
| TLK1 (Bait) | 19 | 17 |
| DSP | 9 | N.C. |
| HSPA4 | 7 | 6 |
| HSPA4L | 7 | 7 |
| RNH1 | 6 | 5 |
| PCNA | 6 | N.C. |

B

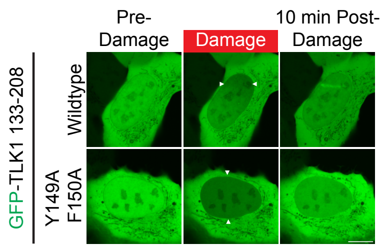


C

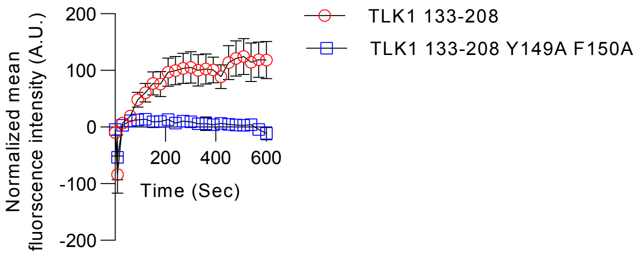
